# MATTE: anti-noise module alignedment for phenotype-gene-related

**DOI:** 10.1101/2022.05.29.493935

**Authors:** Guoxing Cai, Zhan Zhou, Gu Xun

## Abstract

**Purpose:** Although many transcriptome analysis methods find fundamental interactions or markers of some phenotypes, preservation of module or network is still a challenge.

**Methods:** The study developed a method to directly compare the transcriptome data of phenotypes and present the differences modularly, called Module alignedment of TranscripTomE(MATTE).

**Results:** MATTE performs better under high noise than differential co-expression(DC) clustering in the simulation experiments but still detects differential expression(DE) and DC genes. After subsequent annotation of cell types in single-cell data, MATTE obtained the best scores in both supervised and unsupervised learning, i. e. MATTE found meaningful markers. Finally, we apply MATTE in analyzing the transcriptome of Breast Cancer(BRCA). We have found five BRCA subtypes, and the characteristic of one subtype is detected in the form of a module network.

**Conclusion:** MATTE can find meaningful genes and modules, thus facilitating the downstream analysis task to obtain insight into biology.

## Introduction

One of the significant objectives of the multi-omics analysis, including transcriptomics, is to identify genes or recognize related mechanisms associated with phenotypes. With the development of these years, researchers’ attention has gradually shifted from the analysis in a perspective of a single gene to the module or network that can consider more complex relationships. For example, WGCNA [1] is a typical method to identify complex modules and networks from transcriptome data.

One tends to be concerned about which genes, modules, or network substructures result in a change corresponding to a different phenotype. Therefore, it is necessary to solve the problem of whether the constructed module or network will reappear in samples from different phenotypes, that is module’s reproducibility or preservation. Whether a module is preserved or not can be in a biological meanness. In disease, differentiated modules may be of more concern [2, 3], whereas preserved modules in evolution correspond to conservative functions and are essential. Some studies have included more information like ortholog [4–6] in analyzing species to obtain unknown genes’ functions from co-cluster.

However, there are several difficulties:

- The samples often come from different batches or data sources, in which noise is a crucial confounder.
- The results must be easy to interpret and analyze in both preservation and differentiation modules as the application scenarios differ.
- Complex presentation patterns can appear between modules, such as a part overlaps and the core structure of modules retains. Such patterns are different. A method is expected not to ignore these differences.
- A general analysis framework is needed to suit different cases.

Most current studies focused on how to map modules built from different phenotypes samples. Then the following problem is how to define a suitable metric for deciding whether a module is preserved. As summarized in the previous article [7], it can be roughly divided into network-based or cross-tabulation-based. Network-based approaches, however, have failed to address the problem that modules usually show their meaning only if they are preserved or differentiated. Although the cross-tabulation method can analyze the subdivision components of module differences, the analysis results are affected by many factors, such as clustering methods. Furthermore, step errors can be aggregated in the final results of these approaches.

Some studies also want to construct a differential module directly but can only extract one aspect of data, such as building a differential co-expression network—little research composite DE and DC to detect differential modules or networks. For example, COSINE [8] converges DE and DC on one network, but the problem of low computation efficiency appears in large data. This means that filtering is often required before computing can proceed. However, the filtered out genes can involve some functions, and whether such screening is reasonable also needs to be considered.

In this research, motivated by the natural language process where a word in a sentence can be represented by its context, we hypothesize that genes’ expression should be considered in a global view. Relative differential expression(RDE) evaluates direct expression difference and global genes atlas. A gene’s RDE calculates its expression difference to each gene in each phenotype. A similar strategy can also consider co-expression, defined as relative differential co-expression(RDC).

A Module alignedment of TranscripTomE (MATTE) pipeline is designed to cluster and perform module alignedment. MATTE aligneds modules directly by (Fig1b)

1. calculating RDE or RDC transformed data,
2. clustering to assign genes from each phenotype a label,
3. separating genes into preserved and differentiated modules by crosstabulation.

**Figure 1:**
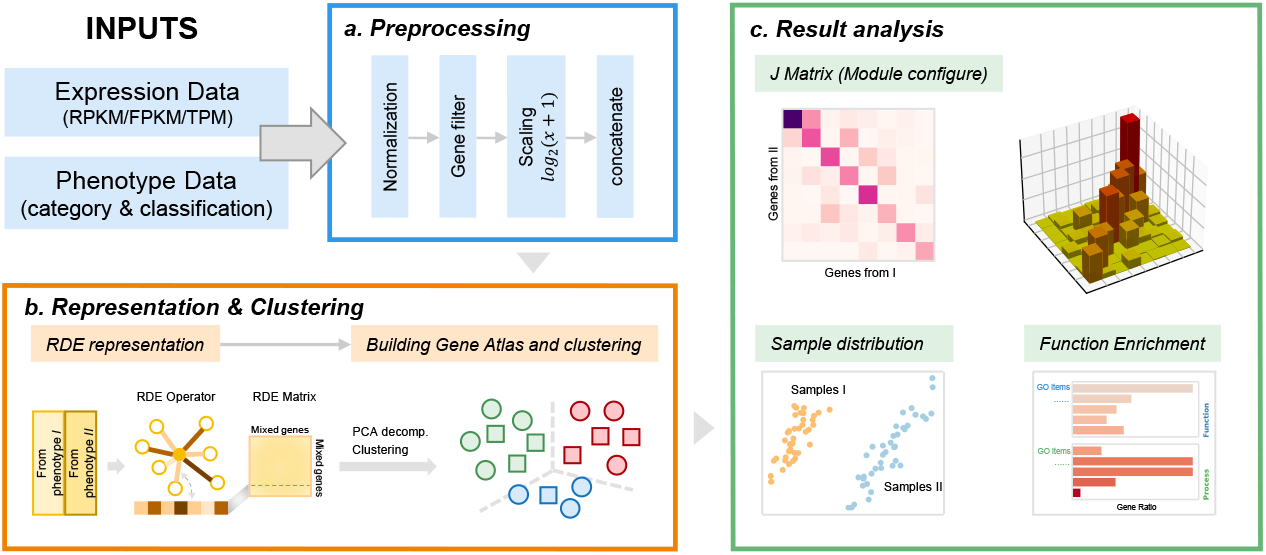
Work flow of MATTE. MATTE takes expression data and phenotype data as inputs. (a):Preprocessing: including normaliztion, filtering, scaling and finaly concatenate data. (b): representation and clustering: using relative difference to characterize genes, decomposition for easy calculations and clustering genes from different phenotypes. (c): Result analysis: relocate genes by their two labels(MC), and analysis specific gene set for function enrichment or sample presentation.

This way, modules can be robustly built for specific phenotypes and aligneded directly.

In the simulation, the advantages of relative difference embedding are demonstrated. MATTE shows a strong anti-noises ability to detect both differential expression(DE) and differential co-expression(DC) compared to other differential clustering methods. Then in single-cell RNA sequencing, transferable markers can be found by a modified MATTE for gene ranking. What is more, MATTE can find meaningful cancer subtypes by modules and explain the subtypes by a module network. These results show MATTE’s robustness, stabilization, and explainability.

## Results

### Overview of MATTE

Most research on transcriptome comparing has been carried out by clustering separately on different phenotypes and aligneding modules [9, 10]. MATTE takes transcriptome data and phenotype data as inputs hoping to construct a space where each gene from different phenotypes is treated as an individual one. By doing so, changes in gene localization from different phenotypes are observed to determine the contribution of genes to phenotypic differences. The same description system for other genes and anti-noise strategy that deals with the potential noise interference of data from different sources is needed to construct such a space.

Here we define the relative differential expression as the difference between each gene and other genes. The noise can be offset as a whole, and global information is delivered to a single gene in relative difference. Such relative differences can be applied to different statistics, such as the mean statistics to measure the degree of differential expression and Pearson correlation coefficients(PCC) to detect co-expression differences. To facilitate the follow-up calculation, we reduce the dimension of the corresponding data by the most common method of principal component analysis(PCA).

By transforming and clustering relative differences, each gene will assign two labels corresponding to two phenotypes. We can relocate genes to module configs(MCs) according to the labels. MC with the same labels is considered not to contribute to phenotype difference(differentiated), while others do(conserved). To evaluate a particular MC’s contribution, signal to noise ratio(SNR) of the eigengene of the MC is calculated. And based on a module’s SNR, we popularize the algorithm for multi-phenotypes gene ranking. And for an MC, gene set analysis can be performed to annotate biological function.

### Simulations show relative difference’s accuracy, robustness, and flexibility

We designed several simulation experiments and compared MATTE with two kinds of methods. Some methods [11–13] aim to identify connection changes of a gene pair. In subsequent experiments, a sum of the distance between a specific gene and other genes is considered its score, and so are RDE and RDC.

The data with differential expression patterns are generated in a multivariate normal distribution. Nine differential expression patterns combine DE and DC in three levels: strong, weak, and none. In the simulation, DE genes are shown as the deviation of the mean value of genes in two conditions. DC genes are presented as a difference in connectivity to other genes in a differential co-expression network, where the PCC builds the graph. All DE and DC genes are labeled as positive and others as negative.

Here, two metrics are used to evaluate methods to find differential expression genes. One is the area under the curve (AUC) of the receiver operating characteristic curve (ROC), which assesses how well the score or rank of genes is consistent with the actual label. The other is the adjusted rand index(ARI) used to test the similarity between the two clustering results.

#### MATTE identified both DE and DC genes accurately

In the following experiments, we try to verify the distance of genes in relative difference space reflects the DE and DC status. As shown in Fig2ab, RDE and RDC achieved the top three AUC in DE and DC, respectively. By fusing the two indexes as RDM, DE, and DC in complex expression mode can be identified; And compared with other methods, it can get the highest AUC, while other methods that claim to be able to recognize both DE and DC have limited effect.

**Figure 2:**
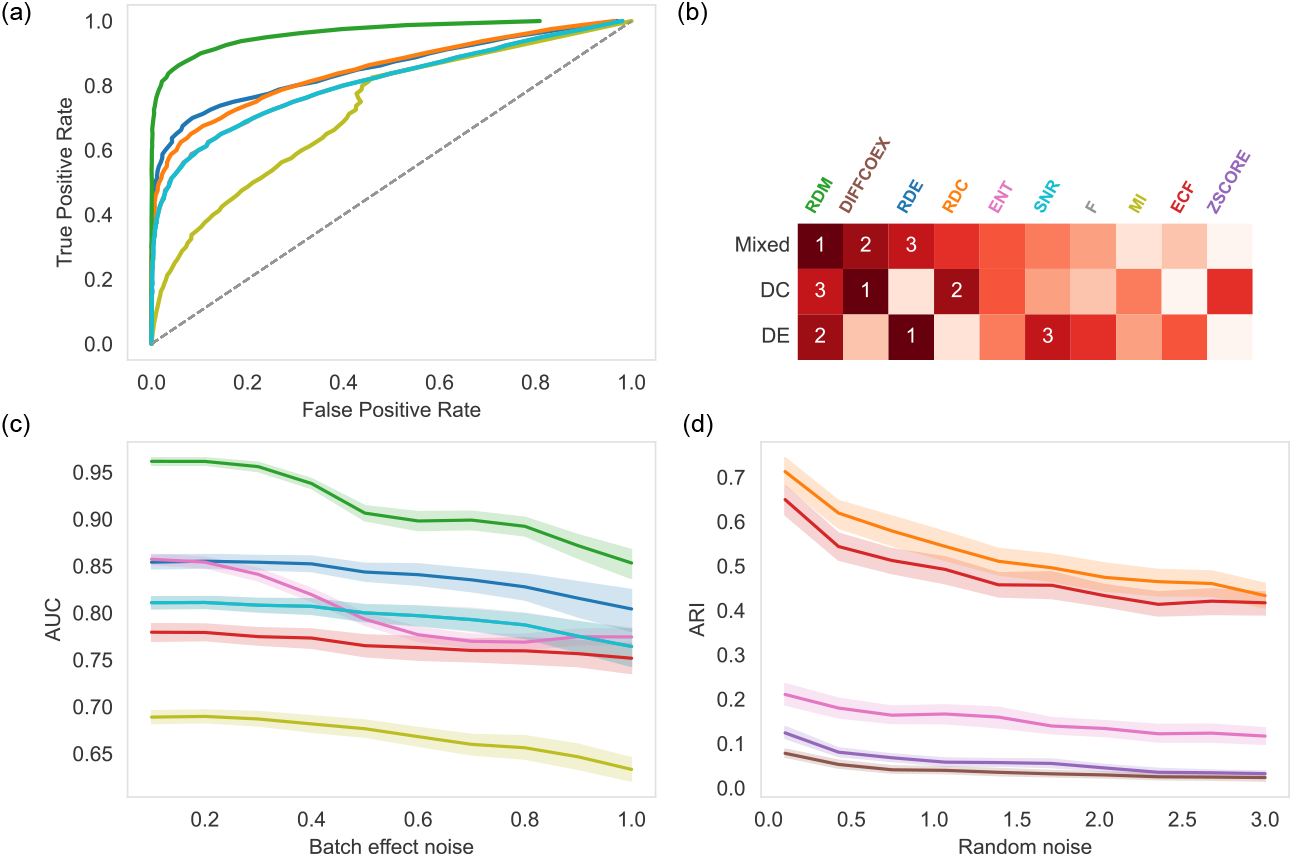
(a): ROC curves of methods in simulations with mixing DE and DC pattern. (b): Rank heatmap of methods in three expression patten simulations. (c): AUC value of ROC when adding batch-effect like noise. (d): ARI of clustering results of before and after adding random noise.

#### MATTE works robustly under a high noise

We test two kinds of noise based on the above experiments, similar to the batch effect and random noise (see method). We randomly assigned a value to 20% of the genes in the generated data for batch-effect-like noise. As shown in Fig 2c, MATTE remains the highest AUC as the noise increases. In the random noise test, we randomly selected 100 genes from real data (100 times, each time the genes were re-extracted). In this test, clustering results using data with and without noise were compared. Compared with the traditional method of calculating the PCC from each edge, the correlation is easily affected by noise, and the RDC is more robust(Fig2d).

### MATTE can identify marker genes in single-cell RNA sequence data

These simulation experiments may not fully reflect the advantages of MATTE, mainly since the differential expression mode is limited to the generation of simulated data. We tested it on the single-cell data. The unique complexity of single-cell data is that many low-expressed genes can not be identified, known as dropouts, because of sequencing accuracy problems. Although some studies have tried to fill the data with deep learning methods, it still needs to consider whether the data will lose its original nature. Identifying cell types is necessary for analyzing single-cell data, and current marker-based single-cell annotation tools rely heavily on prior information [14], that is marker lists. In particular, immune cells can have different cell type annotation depths. As a result, current methods work poorly on it [14, 15]. This part tried to explain the differences in expression in cell types by constructing the module. By building the corresponding modules, the expression of the gene individual can reduce the influence of the dropout event on the cell type annotation. A modified method of MATTE identified markers in the view of the module is shown in Fig3a. In brief, a gene’s score is calculated as the sum of SNR of the module the gene belongs to in each phenotype pair.

**Figure 3:**
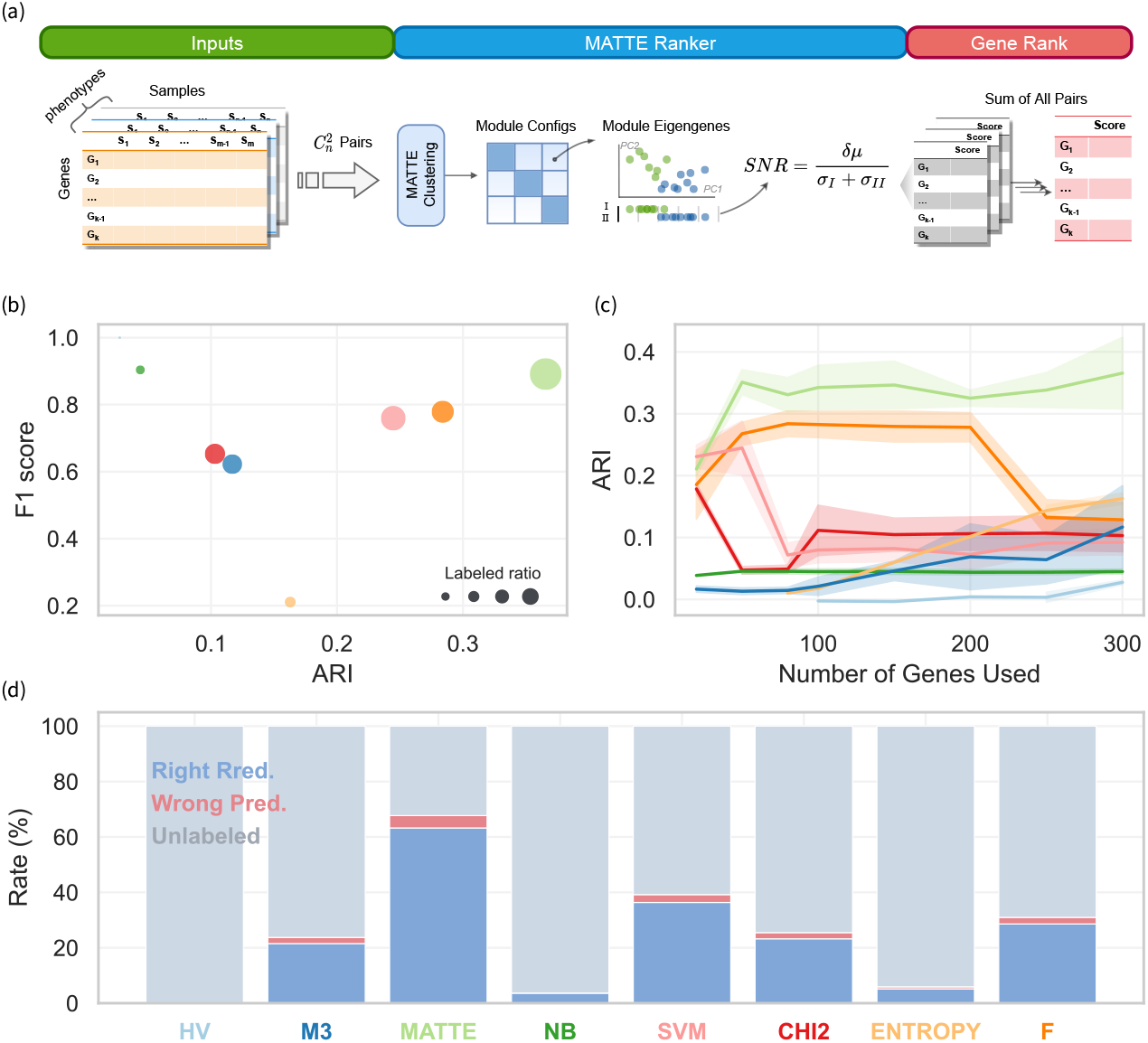
Gene Ranking and results of scRNA seq data.(a) Modified Pipeline of MATTE used in gene ranking: each phenotype pairs will be evaluated separately and in a calculation, SNR of each module will be take as a gene’s score. Results from each turn will be added as final score. (b) Average ratio of prediction of methods: MATTE predict most cells. (c) With the number of marker genes increases, the performance of unsupervised learning task changes.(d) Comprehensive performance of methods in this task.

We applied MATTE, classical feature selection method, differential coexpression method, and differential gene identification method for single-cell data to PUMCBench [16]; the 10Xv2 data set was used as the training set to search for the critical marker genes in six cell types appearing in all data sets. The corresponding rank list is applied to the support vector machine(SVM)-based classifier with a linear kernel function. Based on the recommendations of the previous study [14], we labeled the sample with a prediction probability of less than 0.7 as unknown. And calculate F1 as an evaluation index in the labeled samples. To rule out the influence of the number of the best-selected genes on our test results, 20-300 genes were selected, and the best results were analyzed.

The results showed that MATTE could identify essential genes representing cell types and scored significantly higher on the classification task than other methods(Fig3b). The MATTE classifier had the fewest unlabeled cell types, and the top three overall predicted F1, which predicted the most significant number of cells(Fig3d).

Some methods based on expression variation, such as the depth-adjusted negative binominal model (NB) and Brennecke’s highly variable gene method (HV), have higher F1 values. Still, the ratio of labeled cells is much smaller than MATTE. This may indicate the existence of some distinct cell populations with significant differences in the expression of a few genes.

As the number of selected genes increases, the prediction accuracy of previous classification tasks will also gradually increase because of the low weight of the confounder genes in the SVM classifier. The classification cannot reflect the existence of confounders in the rank. Therefore, we tested it further in an unsupervised task with hierarchical clustering in the euclidean distance. And ARI is used as evaluating indicator. Then, if the features include features that cannot reflect the cell type, the corresponding ARI will decrease. As a result, there are many confounding genes in the marker list identified by most methods; that is, ARI decreases with the increase of the number of used genes. In contrast, the MATTE-sequenced list contained fewer confounding factors. ARI may even rise as the number of genes increases(Fig3c).

### Case study: transcriptome of breast cancer

We further hope to show how to analyze a specific biological case using MATTE. Here we select Breast Cancer(BRCA) in the TCGA project. Breast cancer is one of the most common cancers of women, and the detection of subtypes of BRCA has attracted many studies’ attention [17, 18]. Most used PAM50 [19] subtyping can be used to select an appropriate treatment for patients. In contrast, triplenegative breast cancer(TNBC) lacks therapeutic targets, and drugs are limited, causing the challenge of medicine. In recent years, many studies have focused on finding druggable targets and developing drugs available for TNBC [20, 21]. Here, we hope to compare the normal and cancer transcriptome of breast cancer, understand the potential biological information, and promote the development of relevant, targeted drugs.

We first performed routine MATTE analysis on cancer and normal samples in TCGA. Still, to better mine the data characteristics, we ran the MATTE pipeline of detecting DE and DC, respectively, for cancer data to contain both. The module then characterizes the sample. In short, it is to extract the sample difference information contained in the genes in the module through PCA, that is, module eigengene(ME). The results of the two processes are combined. Subsequently, the module with a low signal-to-noise ratio(SNR) was deleted (< 0.5). Then we use hierarchy clustering and correlation distance to cluster the samples.

After obtaining the subtypes of the sample, we then analyzed the overall survival time. The log-rank test results showed that the p-value was 0.006, which showed significant survival time differences between different subtypes. Specifically, the prognosis of G3 is better, and that of G4 is worse. Later, we compare the corresponding clusters with the PAM50 and find that the G2 and PAM50-Basel are consistent and related. At present, PAM50-Basel is considered synonymous with TNBC in many pieces of research.

We hope to analyze the G2 typing from the perspective of modules to obtain some new conclusions. Then we extract the module network that significantly distinguishes G2 (Fig5c). The genes in the core network were analyzed by functional enrichment. The enrichment analysis can be divided into three parts. Regarding the enrichment of a chromosome location, chr16p13 is exceptionally significant. Many studies have confirmed that mutations and abnormal expression of the region are highly correlated with breast cancer [22–24]. The result of gene enrich in gene ontology(GO) shows the microtubule cytoskeleton involved in mitosis is highly correlated with the development of TNBC, which is highly consistent with the clinical practice and some research [25, 26].

**Figure 4:**
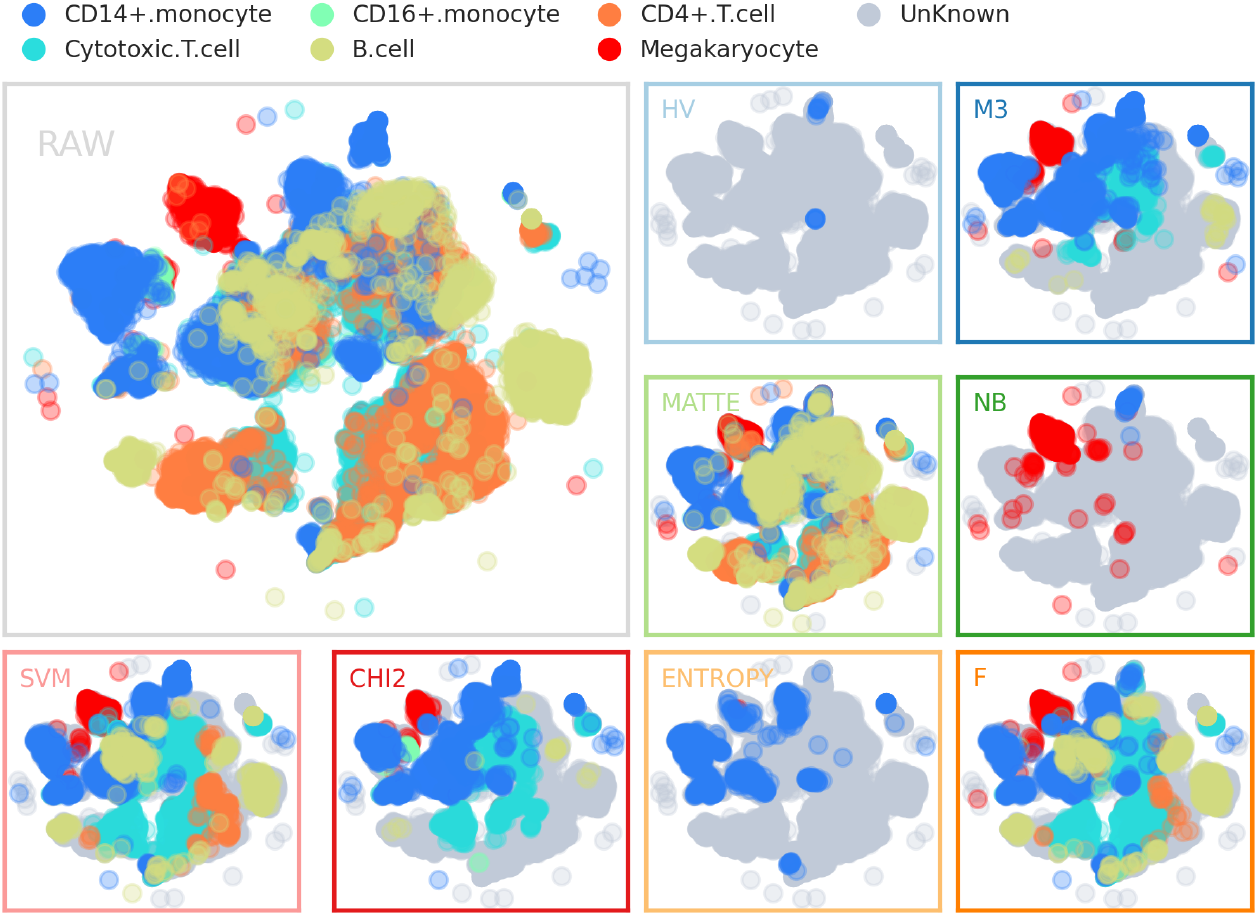
t-SNE Visulization of preditions of methods. Each color refers to a cell type, while grey presents unknown cell types(probability of predition is lower than 0.7).

**Figure 5:**
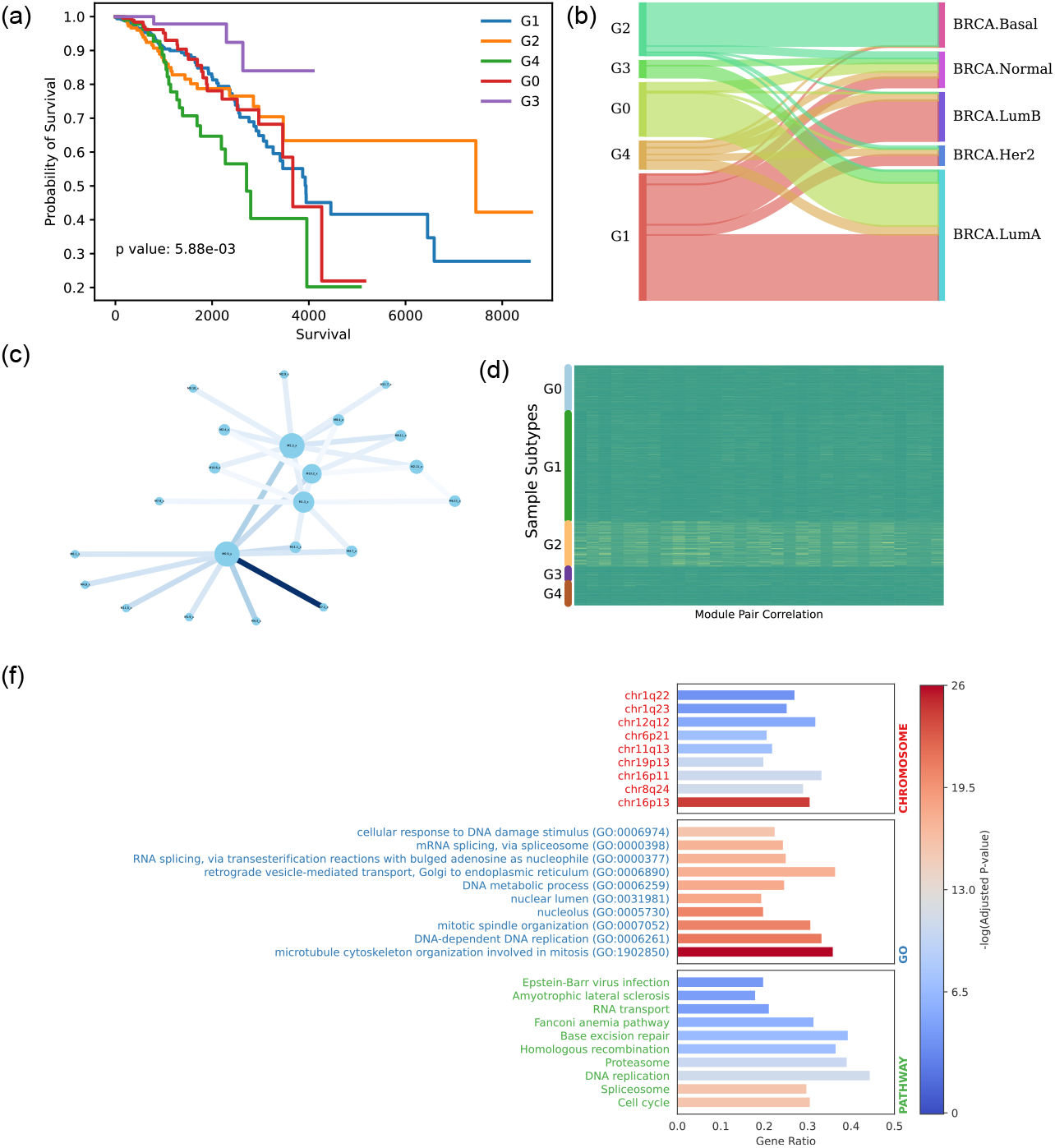
Subtypes of BRCA detected by MATTE.(a) Survivals of five BRCA subtypes, getting p-value 0.006 by multivariate log-rank test. (b) Sanky plot showing the overlaps of our subtypes with PAM50. (c) Core module network that specify G2. The color of edge of the graph represents the difference of G2 and other subtypes. (d) Correlation of pairs of modules in the (c). (f) Function enrich analysis of genes in the core modules network as (c) draws.

## Discuss

Although the perspective of modules has been applied to many studies and methods, aligneding modules has not been considered in recent years. We do not try to analyze and aligned the modules that have been detected but identify the modules directly from the raw data. In this way, it is hoped to reduce the error caused by building modules separately and accumulate in the final result. In our method, the change of genes in the clustering results replaces the specific distance calculation to reduce the impact of random disturbance on the final results. The number of clusters is the scale or resolution of the analysis. We believe that on this issue, based on the theory of causal emergence [27], more information can be obtained than a single gene.

There are all kinds of noise in transcriptome data. Although there are various methods to eliminate the corresponding noise, it is still valuable to construct methods with the ability of anti-noise. The method based on module alignedment also makes MATTE robust to deal with fluctuations. It is conceivable that if noise is generated for the whole, such noise will be eliminated in the alignedment between a single gene and the others, while those genes that produce differences will make the differences more evident compared with context genes.

We hope to introduce network theory and deep learning methods into MATTE in the future. As mentioned above, the expression of a single gene is considered in a broad context, but not all genes can affect gene expression. It is attractive to introduce transformer models, attention strategy, and graph neural networks into this field.

## Materials and Methods

### Overview of MATTE calculation

The core idea of MATTE is to construct a special space in which each gene from each phenotype has a location. By comparing the difference in gene location between the two phenotypes, we can determine whether the gene has an apparent difference between the two phenotypes. This comparison can not be carried out directly in the original data space because of the sample size disparity.

The average value is used in this study to eliminate the influence of sample order. It is worth noting that it is possible to use other statistics to describe the whole, such as the median.

Here, we use a relative differential expression to embed genes. When considering the DE, the difference between each gene and other genes is calculated in absolute value to convert this value into the distance. It is worth noting that using different statistics to describe the whole is possible, such as the median. In the condition of DCE, a normalized spectral embedding is performed based on the graph constructed from Person correlation ecoefficiency.

This representation is consistent with the idea that the distance from three points in a two-dimensional plane can determine the position of an issue. Similarly, in NLP, words are expressed in the context of a phrase.

A new problem arose: a distance in this space can be a differential change. We perform clustering to define the change of location of genes. Compared to setting a threshold of distance, clustering is thought not to overfit and has an ability to anti-noise. Moreover, genes do not work alone; they cooperate. The results in the form of gene sets can make users easy to understand the biological function difference between input phenotypes.

Specifically, genes from different phenotypes are considered independent and clustered together, further distinguished by analyzing changes in the labels of the homogenous gene in different phenotypes. If two labels of a gene corresponding to two phenotypes are the same, the gene is considered not to contribute to the divergences of two phenotypes. We constructed a matrix showing module overlaps for a more detailed analysis, drawing on the cross-tabulation method, where each unit is defined as Module Config (MC). It can be concluded that the differences of each MC lead to phenotypic differences.

### Detailed Steps

#### Data preprocess

Data preprocessing is included by default in the pipeline. The default preprocessing is as follows:

1. Normalization In bulk RNA sequencing data, fragments per kilobase of exon model per million mapped fragments(FPKM) or reads per kilobase of exon per million reads mapped(RPKM) in the data are converted to transcripts per kilobase million(TPM).

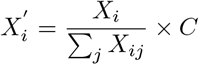 Where 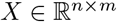 is transcriptome data of a phenotype with *n* genes and *m* samples; *i* is a specific gene, *j* is a particular sample; *C*is a constant. When converted to TPM, $C = 10 ^ 6 $(default). In single-cell data, 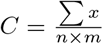.
2. Gene filtering Delete genes with low expression. In this study, genes’ expression lower than 1 in over 90% of samples will be dropped.
3. Scaling Use log transform to reduce the influence of genes with extremely high or low expression.

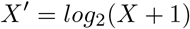

#### Gene Embedding

Gene embedding is based on the difference between a gene and all other genes. For DE, absolute of means’ subtract is used. And use a PCA to decompose the matrix.

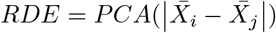

For DC, a network where the nodes are genes and the edge is the PCC between genes is first constructed.

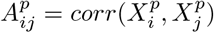

Then perform a spectral embedding where a normalized Laplacian matrix is used to decompose. And finally, concatenate embeddings from two phenotypes.

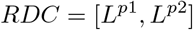

#### Clustering

In this study, the absolute value distance of the PCC is used to cluster the reduced-dimension data with K-Means.

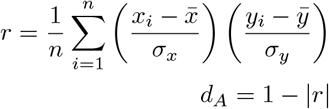

#### Analysis

After clustering, each gene has two labels, *l*_*p*1_ and *l*_*p*2_, corresponding to two phenotypes. By the labels, all genes were further divided into a single MC that contains genes with the same two labels.

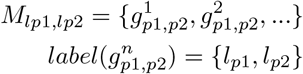

where *label* is a function that maps genes to their labels.

For each MC, module eigengene(ME) is computed to characterize samples in the view of the module. Referring to WGCNA, ME is the first component of data. The data is the expression of genes in the MC when considering DE. While considering DC, the data is the inter-individual correlation(IIC) referring to the previous studies [28, 29].

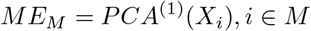

Here we use IIC for most variant gene pairs in the MC, that is:

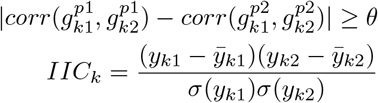

Where *y*_*k*1_ and *y*_*k*2_ are the expressions of genes *g*_*k*1_ and *g*_*k*2_, two genes are from the MC.

### Gene ranking

#### Based on distance

This study calculates the Euclidean distance of two genes corresponding to phenotypes based on gene embedding described previously. To detect both DE and DC genes, we merged distances of RDC and RDE to Relative Differential Merged expression(RDM) based on standardization.

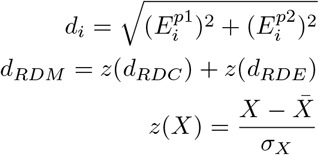

#### Based on module

Based on the MATTE framework, we design a module-based gene rank method. The core idea is to represent the score of genes in the module by the total score of the module. This study uses the SNR to calculate the scores. For the SNR of module *M*:

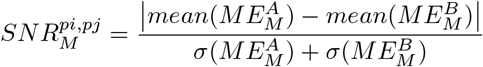

Where 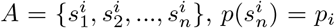 is a set that contains all samples of one phenotype.

When analyzing multiple phenotypes, scores of phenotype pairs are added:

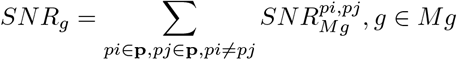

### Simulations

#### Generation of data

The generation of the simulation data is based on the `numpy` package’s multi-variate normal distribution method. We consider genes’ expression pattern as a mixture of two modes: DE and DC. There are nine mode combinations as each mode has strong, weak, and none three levels. The genes with no DE and DC are the negative genes. The method needs to provide a semidefinite matrix as the covariance matrix of the variables. The following describes how the covariance matrix is generated.

It is known that the covariance matrix is the result of the PCC and standard deviation of two variables. Therefore, the covariance matrix can be obtained by generating the PCC matrix. In this study, it is considered that *σ_X_* = *σ_Y_* = 1.

The properties of PCC can be summarized:

1. Is a positive semidefinite matrix
2. Diagonal elements are one.

The covariance matrix can be obtained by generating an arbitrary matrix *E*.

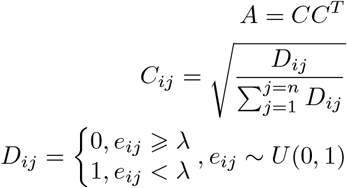

It can be proved that *A* is a positive semidefinite matrix whose diagonal element is 1, the non-diagonal element is less than 1, and is expected to be λ.

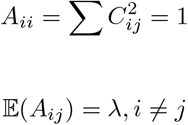

#### Noise simulation

##### Random Noise

In this study, it is considered that the noise conforms to the normal distribution:

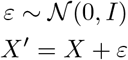

Where *I* represent the variance of the normal distribution of the noise, a different I value is taken to simulate the noise intensity. In this study, value is assumed to simulate the noise intensity.

##### Batch-effects-like noise

We randomly select a certain proportion of genes for the batch-effects-like noise and add a fixed amount of expression to them.

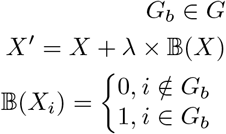

#### Evaluations

##### Classification

In this step, the connectivity of the gene is taken as the score for the gene.

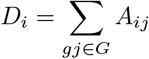

Where the *A* is the weight of the edge, the Area under the Curve(AUC) of the receiver operating characteristic curve(ROC) is used as the ranking index.

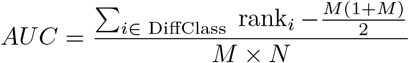

*M* and *N* are the number of positive and negative labels, respectively. *rank* is the result of gene sequencing for a particular method.

##### Clustering

Adjusted Rand Index(ARI) is used as the indicator to compare the labels obtained from the data with and without noise.

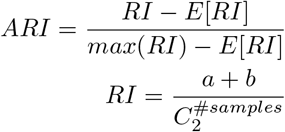

Where *a*, *b* is the number of pairs of elements in the same class and not in the same category in both tags.

### Single-cell RNA sequencing Analysis

One of the challenges of single-cell sequencing data is the dropout of low-expressed genes due to instrument accuracy. A dropout can be thought of as a specific type of noise, making labeling and annotating cells difficult. Some of these methods rely on a priori information that influences the performance decisively. We hope to apply the modified MATTE for gene ranking and other multiple methods to this problem, select several features based on ranking, and annotate the cell type according to the chosen features.

Specifically, the top *k* features of the ranking (*k* ∈ {20, 50, 80, 100, 150, 200, 250, 300}) is used in the following two parts. The first one is a supervised learning task, where a linear kernel support vector machine(SVM) with probability calibration is used to label cell types. Label cells with a maximum prediction probability less than 0.7 as unknown cells and the *F*_1_ score as the indicator.

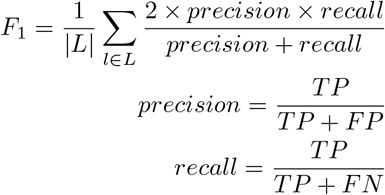

*TP* is the true-positive ratio, *FP* is the false-positive ratio, *FN* is the falsenegative ratio, and *L* is all cell types.

The SVM classifier can recognize important features by assigning the linear kernel’s weight as the number of features increases. The experiment does not reveal whether the gene in front of the sequence contains some of the confounders. Thus, the second part is designed where cells are clustered on K-means using Euclidean distance. Both parts were tested on six cells common to seven data sets of PumcBench. 10xv2 data set is selected as training data and verified on the other six data sets.

### BRCA bulk RNA sequencing Analysis

For DE and DC, the routine process of mate was calculated. Then, we consider that the ME calculated by two results is spliced after the Z transform.

The ME of modules whose SNR is higher than 0.5 is used as the samples’ feature in sample clustering. A hierarchy clustering with PCC distance groups samples into five classes.

Overall survival(OS) analysis is based on python package `lifelines`. A multivariate log-rank test shows if different sample classes have different prognoses.

Subsequently, we compared our BRCA genotyping with PAM50 and found a high overlap between G2 and PAM50-Basel. So we go further and analyze the characteristics of G2. The feature connections of G2 are extracted, and the modular network is constructed. Network visualization is based on the python package `networkx`.

Function enrich the analysis of core network is based on `gseapy`, where gene set data is from `enrichr`.

### Briefings to the compared methods

#### Differential Co-expression related

1. Expected conditional F statistic(ECF) [11] calculates the F statics under the expected condition. ECF combines DE and DC. Python implementation refers to the R package `cosine` [8].

The following three methods’ python implementation refers to R package `dcanr` [30].

2. Z-score [12] converts the PCC of gene pairs into statistics of gene triplets.
3. Entropy [31] of PCC can be calculated based on probabilistic graphical models.
4. DiffCoEX [32] constructs a scale-free network as WGCNA does and uses the topological overlap to calculate distance.

#### Feature ranking methods

1. ANOVA F-value is based on the sum of squares and the ratio of intra-and inter-group deviations in different label groups. In this study, the features are ranked by the value of the F statistic.
2. The chi-square value measures the dependency and independence between data and labels.

#### DE methods for scRNA

M3drop is an R package for single-cell expression data’s DE analysis. Three differential expression methods were used in this study.

1. M3Drop: Under the null hypothesis, it is considered that the dropout ratio and average gene expression follow the Michaelis–Menten equation.
2. The negative binomial model (NB) models each observation as a negative binomial distribution.
3. Brennecke’s highly variable gene method(HVG) looks for genes with significant changes in gene expression. It is considered that there is a linear relationship between the square of the coefficient of variation and the average amount of expression.

### Source code and data availability

BRCA transcriptome can be downloaded from the TCGA project. The single-cell sequencing data set is collated in the previous article [14, 16]. Six cell types that appeared in all data sets were selected for validation and testing from the single-cell data set.

Machine learning methods are based on sklearn [33] and Biopython [34]. The source code of MATTE and case analysis can be found on Github(https://github.com/zjupgx/MATTE). The package can be downloaded and installed via PyPi(https://pypi.org/project/MATTE/) and the MATTE documentation(https://mattedoc.readthedocs.io/en/latest/) is available for users to refer to.

